# An open multimodal spatial resource integrating same-tissue transcriptomics, proteomics, and histology

**DOI:** 10.64898/2026.08.17.742355

**Authors:** Emily Duchini, Colista Tsao, Jason Madore, Thomas Ashhurst, Jeneffer De Almeida Silva, Joo-Shik Shin, Ruta Gupta, Geoffrey McCaughan, Umaimainthan Palendira, Ken Liu, Angela Ferguson, Felix Marsh-Wakefield

**Author notes:** Authors contributed equally (Angela Ferguson, Felix Marsh-Wakefield).

## Abstract

Spatial transcriptomic and proteomic technologies provide complementary insights into tissue organisation, cellular phenotype and function, yet integrating these modalities on the same tissue section remains technically challenging. Sequential workflows must preserve RNA integrity, antigenicity and tissue morphology while maintaining accurate spatial registration. At present, publicly available multimodal datasets suitable for computational method development remain limited.

Here, we present a workflow for sequential 10x Genomics Xenium spatial transcriptomics, COMET cyclic immunofluorescence, and haematoxylin and eosin (H&E) histological staining on the same formalin-fixed paraffin-embedded tissue section. We demonstrate this approach across multiple biologically distinct human tissues, including tonsil, hepatocellular adenoma, and matched tumour and non-tumour hepatocellular carcinoma, illustrating the widespread applicability of the workflow beyond a single tissue type. Following image registration, Xenium-derived cell segmentations were applied to protein images to generate integrated single-cell transcriptomic and proteomic measurements for downstream analyses.

To facilitate community reuse, we publicly release four representative aligned tissue cores together with transcript coordinates, multiplex protein images, H&E images, cell segmentations, and integrated single-cell datasets. We additionally introduce UnumLocalia, an open-source visualisation and data extraction tool that enables interactive exploration of aligned multimodal images, supports user-defined cell segmentation, and allows export of integrated single-cell data for downstream analyses.

Together, this technical protocol, workflow, software, and openly available dataset provide a reusable resource for multimodal spatial biology, supporting advances in biological discovery, computational method development, multimodal data integration, and validation of emerging analytical approaches across complementary spatial technologies.

## Introduction

The tissue microenvironment is a highly complex spatial ecosystem in which coordinated interactions among diverse cell types underpin tissue homeostasis, disease progression, and therapeutic response. Preserving tissue architecture and spatial context is therefore essential for generating biologically interpretable multi-omic data that accurately reflect native cellular states and interactions.

Bulk and single-cell RNA sequencing have transformed our understanding of cellular heterogeneity by enabling quantitative profiling of gene expression across diverse cell populations. However, these approaches typically require tissue dissociation, disrupting cell–cell interactions, inducing stress-related transcriptional artefacts, and eliminating spatial information critical for understanding tissue organisation and function^1,2^. To address these limitations, spatial transcriptomic technologies were developed to localise mRNA transcripts within intact tissue sections, enabling spatially resolved transcriptomics, and in some platforms, single-cell or subcellular spatial transcriptomic profiling^3^.

Despite these advances, transcriptomic measurements alone are often insufficient to infer cell phenotype and function. mRNA abundance does not consistently correlate with protein expression due to extensive regulation at post-transcriptional and post-translational levels, differences in protein stability, and spatially constrained protein localisation^4,5^. In spatial contexts, this disconnect can lead to inaccurate assumptions about functional cell states if transcriptomic data are interpreted in isolation. Accordingly, there is growing interest in integrating spatial transcriptomic and spatial proteomic measurements to improve cell identification, functional interpretation and biological insight^6^.

In parallel, spatial proteomic technologies have advanced rapidly, enabling increasingly deep and comprehensive characterisation of protein expression within intact tissues. Recent work has demonstrated the feasibility of profiling hundreds of proteins by multiplex immunostaining, including the simultaneous detection of more than 580 proteins in human head and neck squamous cell carcinoma, alongside spatial transcriptomic mapping of approximately 18,000 mRNA targets using a whole transcriptome atlas^7^. These advances highlight the growing potential of combining high-resolution spatial transcriptomics with high-plex spatial proteomics to achieve a more complete functional view of tissue biology. Furthermore, it is demonstrated that this is possible despite the increased technical complexity of maintaining RNA integrity, antigenicity, tissue morphology, and imaging quality across sequential assays. Reflecting this progress, technological developments have begun to enable the combination of multiple spatial assays on a single tissue section. Multiplex immunofluorescence has been combined with haematoxylin and eosin (H&E) staining and imaging mass cytometry on formalin-fixed paraffin-embedded (FFPE) human tonsil tissue, demonstrating compatibility between protein-based and morphology-based imaging modalities^8^. More recently, spatial transcriptomics using the 10x Genomics Xenium platform has been followed by COMET cyclic immunofluorescence (cyclicIF) and H&E on FFPE lung carcinoma sections, illustrating the feasibility of integrating high-resolution transcriptomic and proteomic measurements on the same physical tissue section^9^. However, performing multiple spatial assays on the same tissue section remains technically challenging. Sequential workflows must carefully balance RNA preservation, antigen integrity, spatial registration accuracy, and maintain tissue morphology. Due to these complexities, most existing studies focus on individual tissue types or proof-of-concept demonstrations, limiting their utility for broader method development, validation, and cross-platform benchmarking.

In this study, we present a robust workflow for sequential Xenium spatial transcriptomics, COMET cyclicIF and H&E on the same FFPE tissue section. We apply this workflow across multiple biologically distinct human tissues, including human tonsil, hepatocellular adenoma (HCA), and matched tumour and adjacent non-tumour tissue from a patient with hepatocellular carcinoma (HCC), illustrating its broad applicability, beyond a single tissue type or disease setting. To facilitate reuse of these data, we publicly release four representative aligned tissue cores together with the underlying transcript coordinates, protein measurements, and imaging data. We additionally introduce UnumLocalia, an open-source visualisation and data extraction tool that enables interactive exploration of aligned multimodal datasets, supports user-defined cell segmentation, and exports integrated single-cell transcriptomic and proteomic measurements for downstream analysis. Together, the workflow, software and publicly available datasets provide a valuable community resource for multimodal spatial biology, facilitating biological advances, computational method development, multimodal data integration, validation of emerging analytical approaches, and reproducible reuse of aligned spatial datasets.

## Methods

### Sample collection

Ethics was obtained from the Sydney Local Health District Ethics Review Committee (HREC X20-0552 & 2020/ETH02958). FFPE human tissue samples were obtained from the Department of Gastroenterology and Hepatology, Department of Tissue Pathology and Diagnostic Oncology, and Liver Biobank at Royal Prince Alfred Hospital to be made into a tissue microarray (TMA). Four 1 mm^2^ cores from the TMA were examined in this study, including one tonsil sample, one liver sample obtained from a patient with HCA, and two matched liver samples (non-tumour and tumour regions) obtained from a patient with HCC.

### Xenium spatial transcriptomics

Samples were prepared using the Xenium Prime 5K Human Pan Tissues and Pathways Assay Kit (10x Genomics, PN-1000671) following manufacturer protocols (User Guides CG000578 and CG000760). The TMA block containing FFPE tissue samples was sectioned at 5 μm onto a Xenium Slide, performed by the Charles Perkins Centre Histology Facility (The University of Sydney, Australia). The slide was dried at room temperature (RT) before being placed in an oven at 42℃ for 3 hours, then stored in a desiccator overnight at RT. Deparaffinisation of the tissue section were performed in a thermal cycler (Bio-Rad) at 60℃ for 30 minutes, after which the slide was left to cool down to RT for 7 minutes. The slide was immersed twice in xylenes (Sigma Aldrich) for 10 minutes each, then rehydrated in a graded series of ethanol (Sigma) solutions for 3 minutes each at the following concentrations: 100%, 100%, 96%, 96%, 70%. The slide was washed in Nuclease-free water (Thermo Fisher Scientific) for 20 seconds prior to being placed into a Xenium Cassette then washed with 500μL of phosphate-buffered saline (PBS). 500μL of Decrosslinking Buffer (10x Genomics) was applied to the slide before being placed in a thermal cycler for 30 minutes at 80℃ to release the sequestered RNA from the tissue. The slide was then washed four times in 500µL of PBS-Tween (PBS-T) for 1 minute each.

After deparaffinisation and decrosslinking was complete, the section underwent tissue hybridisation, ligation, amplification and autofluorescence quenching. First, 150μL of Priming Hybridization Mix containing priming oligos (10x Genomics) was applied to the tissue to hybridise to target RNA. The slide was placed in the thermal cycler for 1.5 hours at 50℃, followed by three washes in PBS-T. Next, 500µL of Post-Priming Wash Buffer (10x Genomics) was applied to the slide for a 30-minute incubation at 50℃, followed by three washes in PBS-T to remove unbound priming oligos. 500µL of RNase Mix (10x Genomics) was then applied to the slide for a 20-minute incubation at 37℃ to cleave the RNA strands precisely where the priming oligos are bound. This was followed by three washes in 0.5X saline-sodium citrate buffer containing Tween-20 (SSC-T). 500µL of Polishing Reaction Mix (10x Genomics) was applied to the slide for a 1-hour incubation at 37℃ to standardise the cut RNA and create consistent binding sites for the probes. After three washes in PBS-T, 150µL of Probe Hybridization Mix containing DNA probes (10X Genomics) was applied to the slide to hybridise to the target RNA. The slide was placed in the thermal cycler for an overnight (16–24 hour) incubation at 50℃.

The next day, the slide was washed three times in in PBS-T before 500µL of Post Hybridization Wash Buffer (10x Genomics) was applied to the slide for a 15-minute incubation at 35℃. Following three washes in PBS-T to remove unbound probes, 500µL of Ligation Mix (10x Genomics) was applied to the slide for a 30-minute incubation at 42℃ to join adjacent probe ends bound to the RNA. In doing so, this ligation step forms circular DNA probes that confer high specificity for the target regions. After three washes in PBS-T, 500µL of Amplification Enhancement Master Mix (10x Genomics) was applied to the slide for a 2-hour incubation at 4℃ to prepare the circular DNA probes for efficient amplification. This was followed by a wash in 500µL of Amplification Enhancer Wash Buffer (10x Genomics) for 1 minute, after which 500µL of Amplification Master Mix was applied to the slide for a 1.5-hour incubation at 30℃ to induce rolling circle amplification and generate hundreds of gene-specific barcode copies. After three washes in 1X Tris-EDTA Buffer (Fisher Scientific), the slide was washed in a series of ethanol solutions (70%, 100%, 100%, 70%) for 2 minutes each to prepare the tissue for cell segmentation antibody staining. Following a wash in PBS-T, 500µL of Diluted Xenium Block and Stain Buffer (10x Genomics) was applied to the slide for a 1-hour incubation at RT to minimise non-specific binding in the subsequent antibody staining step. 100µL of Xenium Multi-Tissue Stain Mix containing cell surface and interior antibodies (10x Genomics) was then applied to the slide for an overnight (16 – 24 hour) incubation at 4℃ for downstream cell segmentation.

Following the overnight incubation, the slide was washed three times in PBS-T to remove excess antibodies. 500µL of Xenium Stain Enhancer (10x Genomics) was then applied to the slide for a 20-minute incubation at RT, followed by three PBS-T washes. 500µL of Diluted Reducing Agent B (10x Genomics) was applied to the slide for a 10-minute incubation at RT, followed by three washes in a series of ethanol solutions (70%, 100% and 100%) for 1 minute each. 500µL of Autofluorescence Solution (10x Genomics) was applied to the slide for a 10-minute incubation in the dark at RT to quench unwanted autofluorescence signal, followed by three washes in 100% ethanol for 2 minutes each. The slide was then dried in the thermal cycler for 5 minutes at 37℃, followed by a wash in PBS to rehydrate the tissue then a final wash in PBS-T in the dark. 500µL of Nuclei Staining Buffer containing DAPI (10x Genomics) was applied to the slide for a 1-minute incubation at RT in the dark, followed by three washes in PBS-T. 1mL of PBS-T was applied to the slide before loading it into the Xenium Analyzer for automated cyclic imaging and decoding.

### Post-Xenium tissue preparation

Following acquisition on the Xenium Analyzer, the processed Xenium slide was prepared for sequential COMET staining. Firstly, the tissue section underwent antigen retrieval by submerging the slide in a container filled with pH 9.0 antigen retrieval buffer (10 mM Tris Base, 1 mM EDTA solution containing 0.05% Tween-20, adjusted to pH 9.0) before heating in a pressure cooker at 95°C for 1 hour. After cooling to RT, the slide was washed in tris-buffered saline with Tween-20 (0.1 M Tris HCl, 0.15 M NaCl solution containing 0.05% Teen-20, adjusted to pH 7.5) for 2 minutes. The tissue section was then blocked to minimise non-specific binding by firstly incubating with avidin/biotin blocking kit (Life Technologies) for 10 minutes each at RT, followed by an incubation with horse serum (Life Technologies, 20% in DPBS) for 45 minutes at RT. Lastly, the tissue was blocked with 1X Antibody Diluent/Blocking Buffer (Akoya Biosciences) for 45 minutes at 37°C.

### COMET spatial proteomics

A 15-plex previously optimised primary antibody panel was used to perform cyclicIF using the COMET^TM^ system (Lunaphore) on the processed Xenium slide. The primary antibodies were detected by either anti-mouse, anti-rat or anti-sheep fluorescently-conjugated secondary antibodies (**Table 1**). The antibodies and staining buffers were prepared and loaded into the COMET instrument, along with the tissue section, according to the protocol generated by the COMET Control Software for automated cyclic protein staining and imaging.

Each cycle involved primary antibody staining for 4 minutes at 37℃ followed by staining with secondary antibodies for 2 minutes at 37℃. Imaging was then performed using a combination of the following channels at the described exposure times: DAPI 25 milliseconds, TRITC 250 milliseconds, Cy5 400 milliseconds and FITC 500 milliseconds. After imaging, elution was performed for 2 minutes at 37℃ using Elution Buffer (Lunaphore), followed by quenching for 30 seconds at 37℃ using Quenching Buffer (Lunaphore) to ensure all antibodies were properly removed before the next cycle. The acquired images (magnification, 20x) were automatically stitched and aligned by the COMET^TM^ system. Multi-channel OME.TIFF images with background fluorescence subtracted per channel were exported using HORIZON^TM^ software (Lunaphore).

### H&E staining

H&E staining was performed at the Charles Perkins Centre Histology Facility (The University of Sydney, Australia) as previously described^10^.

### Publicly available multimodal dataset and software

To facilitate reuse of these data, we provide the aligned Xenium, COMET and H&E images together with the underlying transcript coordinates, cell segmentations and protein measurements for all four representative tissue cores (https://zenodo.org/records/21713660). Accompanying these data, we developed UnumLocalia, an open-source toolkit for exploring aligned multimodal spatial datasets (https://github.com/Felixillion/UnumLocalia). Image registration was performed externally in Xenium Explorer (v4.1.1), after which UnumLocalia directly consumes the resulting transformation matrices to visualise aligned transcriptomic, proteomic and histological images without requiring users to repeat image registration. An included clustering script is available for visualising single-cell data (as shown in **Figure 3** and **Supplementary Figure 2**). The software enables interactive visualisation of gene transcripts, protein markers and histological images, supports import of alternative cell segmentations, and provides workflows for single-cell quantification of transcripts and proteins. Quantification tables, images, and clustering outputs can be exported for downstream analysis. Users can also incorporate their own multimodal spatial datasets by supplying image data, molecular measurements and alignment information. Together, the publicly available dataset and software provide a resource for exploring and benchmarking multimodal spatial biology data.

## Results

### Sequential Xenium, COMET and H&E staining can be performed on the same tissue section

We developed a workflow that enables sequential spatial transcriptomic (Xenium), multiplex protein (COMET) and H&E imaging from the same FFPE tissue section (**Figure 1**). Four representative tissue cores (human tonsil, HCA, and matched tumour and non-tumour HCC) were processed using this workflow without obvious loss of tissue integrity or image quality (**Supplementary Figure 1**), demonstrating compatibility between all three modalities.

**Figure 1:**
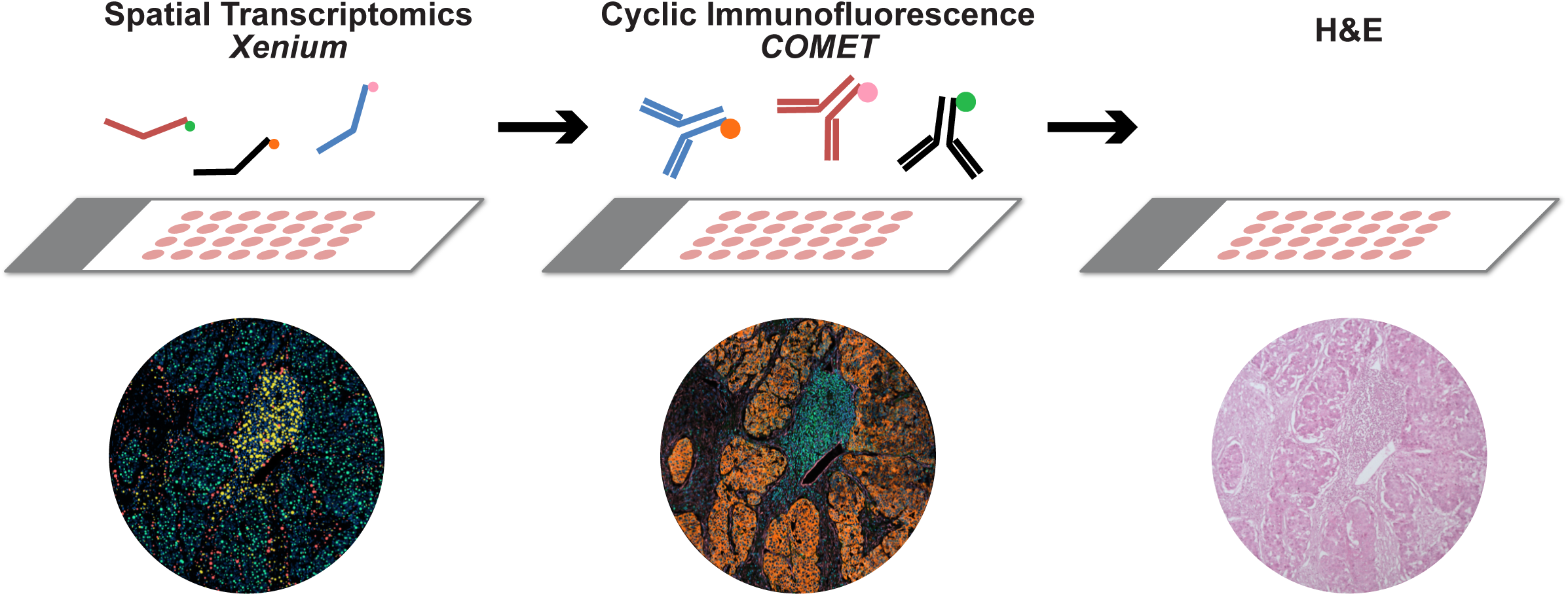
Overview of same-tissue, sequential Xenium, COMET, and H&E staining workflow. One FFPE tissue section firstly underwent Xenium spatial transcriptomics to map 5,001 mRNA targets across multiple biologically distinct human tissues. After the run was complete, COMET cyclic immunofluorescence measuring 15 proteins was performed on the same tissue section, followed by H&E staining.

### Same-tissue gene and protein assessment captures complementary biological information

Following image registration, transcript and protein signals could be directly compared on the same tissue section (**Figure 2**). Examples were selected to illustrate the varying relationship expected between corresponding RNA transcript and protein measurements on individual cells and within tissue architecture. CD20 protein and *MS4A1* RNA demonstrated strong spatial concordance across all four tissues (**Figure 2A**), whereas HepPar1 protein showed minimal agreement with *CPS1* transcripts despite both being hepatocyte-associated markers (**Figure 2B**). *VEGFA* transcripts and VEGFA protein exhibited distinct but biologically relevant spatial distributions for RNA message, protein production, and cellular expression (**Figure 2C**), with transcripts detected in cells and protein extending into the surrounding microenvironment. These examples illustrate that transcript and protein abundance are not necessarily co-localised at the single-cell level, highlighting the importance of integrating complementary molecular measurements from the same tissue section rather than inferring protein abundance directly from transcriptomic data.

**Figure 2:**
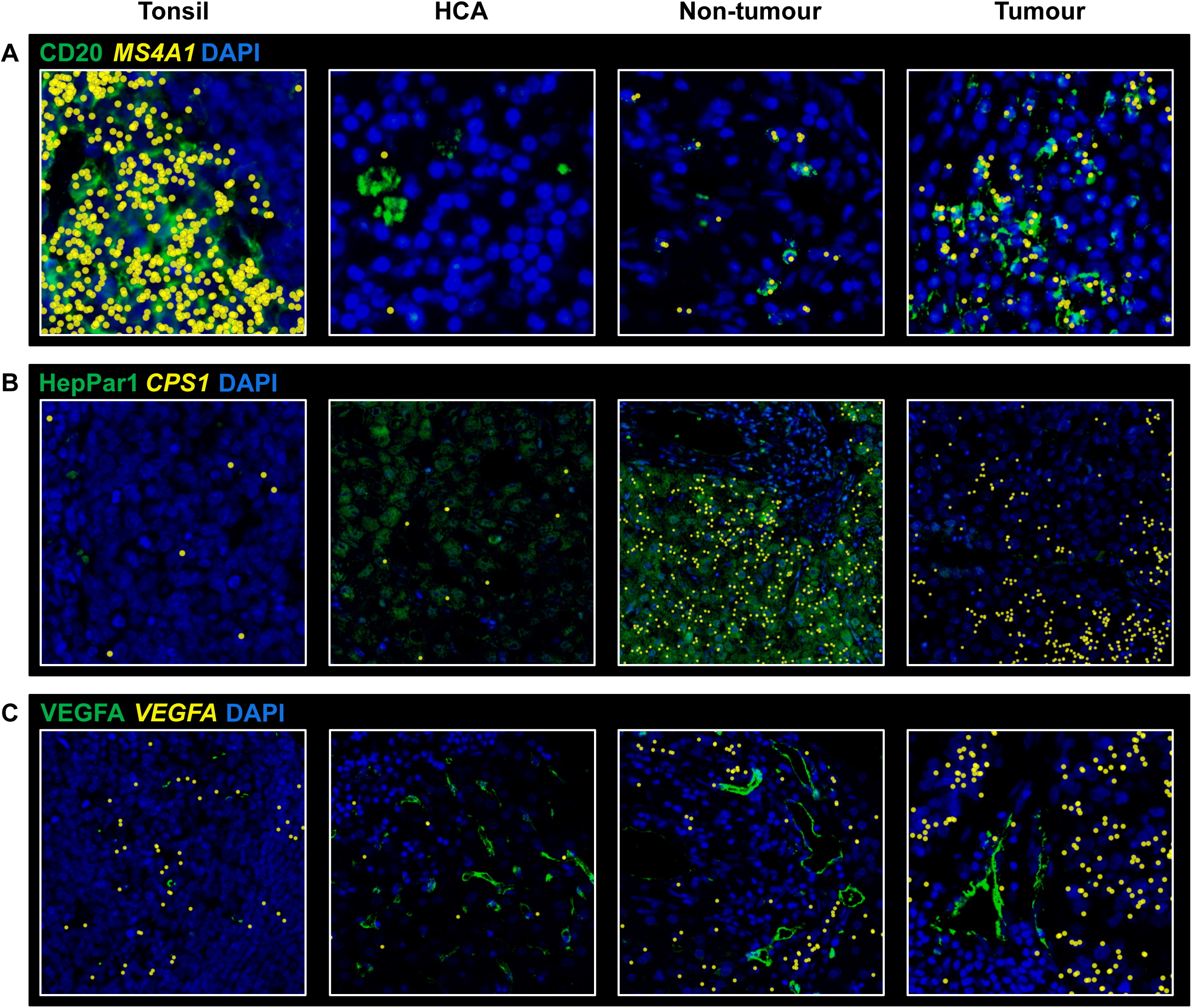
Aligned tissue images of protein and transcript enable direct investigation of complementary molecular features. The expression of matched protein (green) and transcript (yellow) at single-cell resolution were compared across four biologically distinct human tissues, including tonsil, hepatocellular adenoma (HCA), hepatocellular carcinoma (HCC) non-tumour and HCC tumour (left to right). Matched protein and transcript marker pairs presented include A) CD20/*MS4A1*, B) HepPar1/*CPS1*, and C) VEGFA/*VEGFA*.

### Integrated gene and protein measurements produce detailed and complementary single-cell data

Xenium-derived cell segmentations were applied to the aligned COMET images to quantify protein expression for individual cells. Gene-only, protein-only and combined transcriptomic-proteomic datasets were analysed independently using identical single-cell clustering workflows (**Figure 3**; **Supplementary Figure 2**). Agreement between clustering results was quantified using the Adjusted Rand Index (ARI), which measures similarity between cluster assignments while accounting for agreement expected by chance, and the Normalised Mutual Information (NMI), which quantifies the amount of shared information between clustering solutions. Across the four tissue types, combined clustering showed varying levels of agreement with both gene-only and protein-only analyses, demonstrating that each modality contributes distinct yet complementary information to cell-state identification. These findings demonstrate that transcriptomic and proteomic measurements capture complementary aspects of cellular identity within the same physical cells, illustrating the value of truly integrated same-section datasets and providing a resource for the development and evaluation of multimodal integration methods.

**Figure 3:**
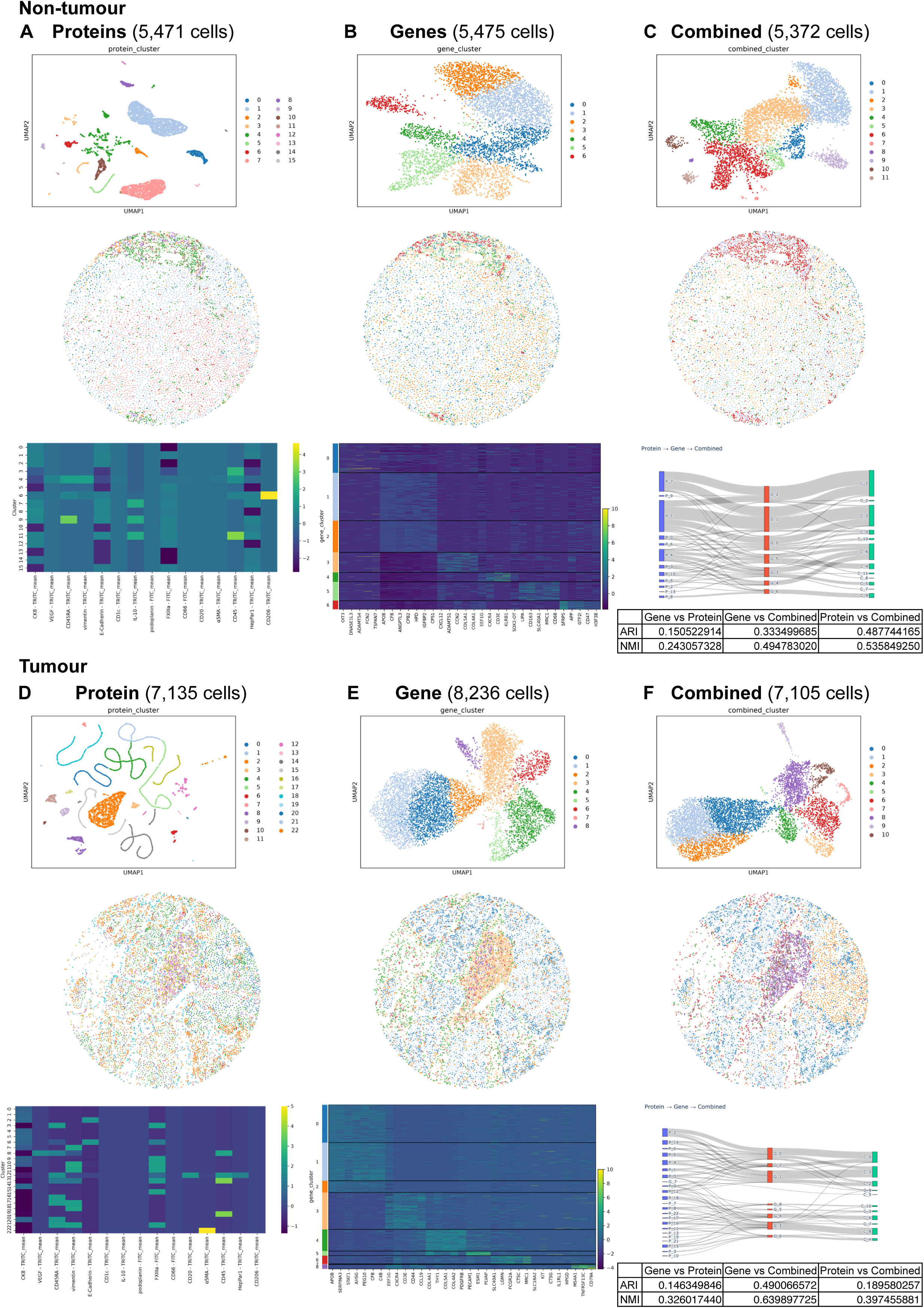
Multimodal integration produces complementary data to identify single cells in tumour and non-tumour samples. Single cells were assessed for protein and gene expression in A-C) hepatocellular carcinoma (HCC) non-tumour and D-F) HCC tumour tissue. Top row shows low-dimensional embeddings computed from (A, D) protein expression, (B, E) gene expression, and (C, F) combined gene and protein expression. Middle row shows spatial maps of cluster assignments from top row. Bottom row shows marker heatmaps of the protein-only and gene-only clustering results. Bottom row right-inset shows mapping of cluster identities across modalities (Sankey) and quantitative agreement metrics (ARI, NMI). Supplementary Figure 2 shows the same panels for the remaining cores (tonsil and hepatocellular adenoma). ARI, Adjusted Rand Index; NMI, Normalised Mutual Information.

### Preservation of tissue landscape is histologically confirmed across all imaging modalities

H&E images were aligned with Xenium transcriptomic and COMET protein images, enabling direct overlay of tissue morphology with molecular measurements (**Figure 4**). Representative examples demonstrate preservation of major histological structures across all modalities, including lymphoid follicles in tonsil and hepatic architecture within non-tumour and tumour liver tissue. These aligned datasets enable molecular measurements to be interpreted within their histopathological context and provide a valuable resource for developing and benchmarking image registration, segmentation, multimodal analysis methods and in gaining biological insight.

**Figure 4:**
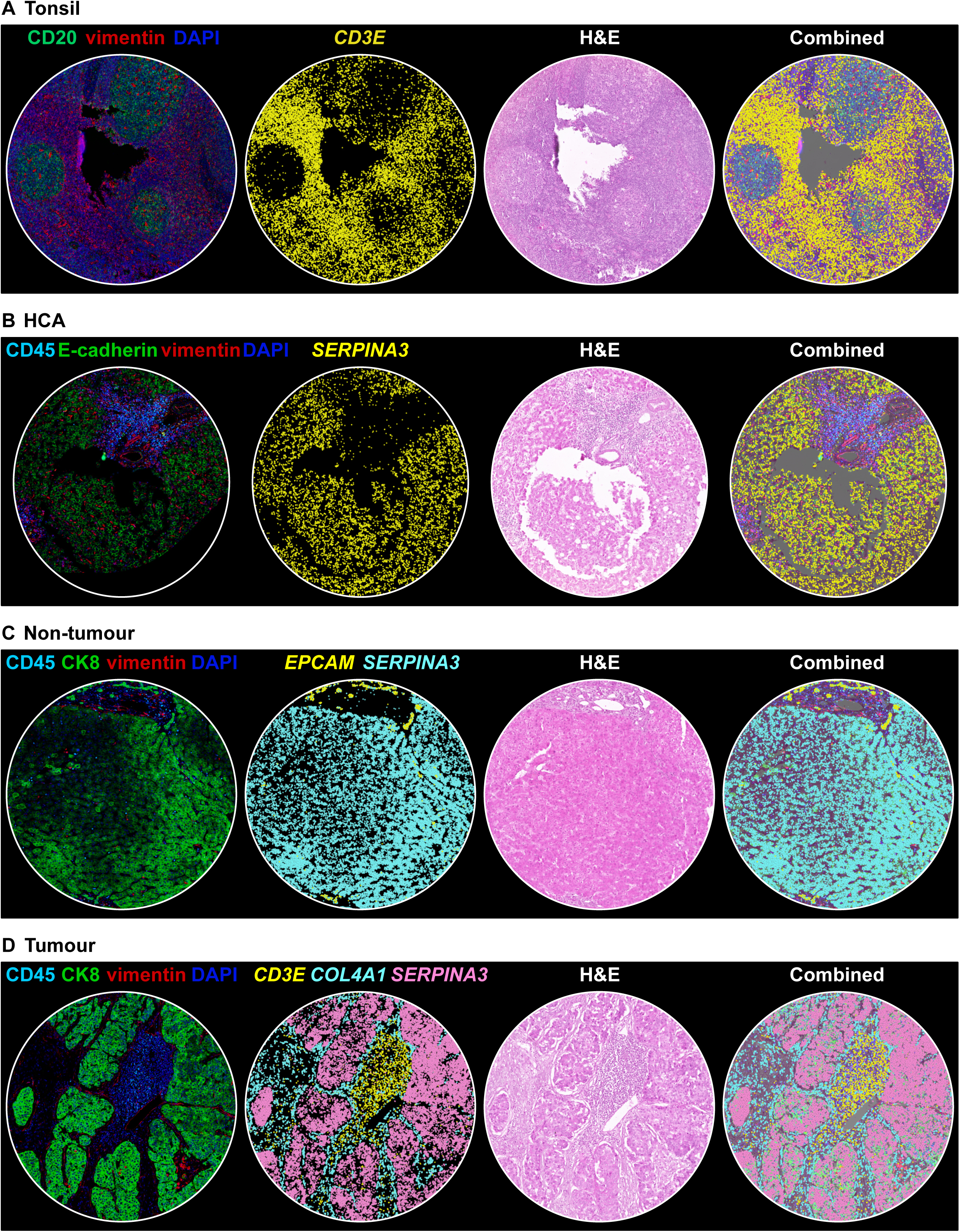
Preservation of biological structures across H&E, transcriptomics and multiplex protein imaging. Representative images of (left-right) COMET cyclicIF, Xenium spatial transcriptomics, H&E staining, and a combined modalities image captured from the same tissue section across four biologically distinct human tissues. A) Tonsil sample, expression of the proteins CD20 (green) and vimentin (red) along with DAPI (blue), captured by COMET, shown alongside the expression of the transcript *CD3E* (yellow) measured by Xenium. B) Hepatocellular adenoma (HCA) sample, expression of the proteins CD45 (cyan), E-cadherin (green) and vimentin (red) along with DAPI (blue), captured by COMET, shown alongside the expression of the transcript *SERPINA3* (yellow) measured by Xenium. C) Hepatocellular carcinoma (HCC) non-tumour sample, expression of the proteins CD45 (cyan), CK8 (green) and vimentin (red) along with DAPI (blue), captured by COMET, shown alongside the expression of the transcripts *EPCAM* (yellow) and *SERPINA3* (pink) measured by Xenium. D) HCC tumour core sample, expression of the proteins CD45 (cyan), CK8 (green) and vimentin (red) along with DAPI (blue), captured by COMET, shown alongside the expression of the transcripts *CD3E* (yellow), *COL4A1* (cyan) and *SERPINA3* (pink) measured by Xenium.

### UnumLocalia facilitates exploration and straightforward utilisation of aligned multimodal spatial datasets

To improve accessibility and utility of the multimodal dataset, we developed UnumLocalia, an open-source visualisation and data extraction tool for aligned spatial transcriptomic, spatial proteomic and H&E imaging data (**Figure 5**). UnumLocalia enables interactive exploration of pre-aligned tissue images, visualisation of selected transcripts and proteins, and application of default Xenium or user-defined cell segmentations. Protein expression can then be quantified using the supplied cell masks and integrated with Xenium transcript counts to generate single-cell AnnData objects for downstream analyses, including clustering and dimensionality reduction. By supporting alternative segmentation strategies while maintaining a common underlying multimodal dataset, UnumLocalia provides a flexible platform for method development, validation and reproducible reuse of the publicly available resource.

**Figure 5:**
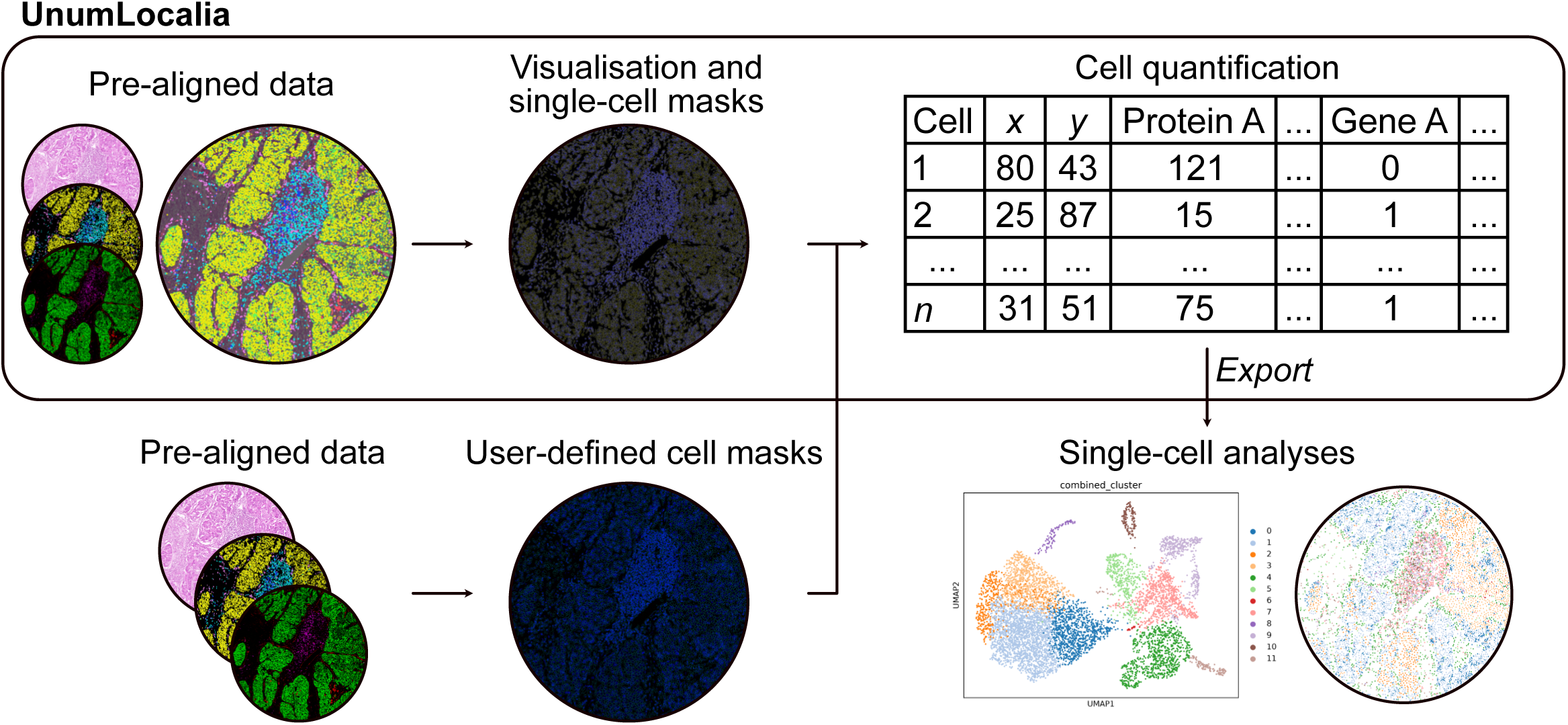
UnumLocalia facilitates exploration and straightforward utilisation of aligned multimodal spatial datasets. Overview of UnumLocalia for visualising pre-aligned, multimodal data (including H&E, mRNA transcript, and protein). Cell masks can be imported to quantify protein and gene expression at a single-cell level. Data can then be exported for downstream analyses.

## Discussion

Spatial technologies are enabling increasingly comprehensive molecular characterisation of tissues while preserving their native architecture. As these technologies continue to mature, there is growing interest in combining complementary modalities on the same tissue section to provide simultaneous transcriptomic, proteomic and histological information. In this study, we establish a reproducible workflow for sequential Xenium spatial transcriptomics, COMET cyclicIF and H&E staining on the same FFPE tissue section. We further provide publicly available aligned high-dimensional molecular datasets from multiple human tissues along with accompanying software to facilitate method development, benchmarking and exploration of multimodal spatial data.

Our workflow builds upon recent studies demonstrating the feasibility and value of combining multiple spatial imaging modalities on a single tissue section. Previous work has shown compatibility between multiplex immunofluorescence, imaging mass cytometry and H&E staining on FFPE tissue^8,11^, while Tran *et al.* (2025) recently demonstrated sequential Xenium, COMET and H&E imaging in lung carcinoma^9^. In parallel, Tan *et al.* (2025) highlighted the rapidly expanding capability of spatial proteomics by combining whole-transcriptome spatial profiling with detection of more than 580 proteins within the same head and neck squamous cell carcinoma tumour specimens^7^. Whereas previous studies primarily demonstrated the technical feasibility of same-section multimodal imaging, the present study extends this concept by providing a reproducible workflow, openly available aligned datasets spanning multiple tissue types, and accompanying software that together create a reusable community resource for computational method development and multimodal spatial analysis.

One of the principal advantages of performing multiple assays on the same physical tissue section is the elimination of uncertainty introduced when integrating serial sections. Although adjacent sections generally preserve overall tissue morphology, individual cells are frequently absent, displaced or sectioned differently, particularly in heterogeneous tissues or regions containing infiltrating immune cells. Same-section imaging enables direct integration and comparison of RNA transcripts, proteins, and histopathological features originating from individual cells and tissue structures, thereby providing valuable unique ground data for benchmarking registration, segmentation, multimodal integration methods, and biological investigations. Importantly, our results also illustrate that transcript and protein measurements should not necessarily be expected to demonstrate one-to-one correspondence. Some markers, such as CD20, showed close correspondence between transcript and protein localisation, whereas others, including VEGFA, demonstrated distinct spatial distributions. These observations highlight that transcriptomic and proteomic measurements can provide either concordant or complementary information depending on the underlying biology, reinforcing the value of integrating both modalities rather than relying on either alone. Differences between mRNA abundance and protein localisation reflect biological regulation and active biological processes as well as technical differences between measurement modalities, emphasising that these datasets provide complementary information adding extra dimensionality to aid biological interpretation.

The availability of pre-aligned multimodal human tissue data is particularly valuable for computational method development. Current challenges in spatial biology include image registration, cell segmentation, multimodal integration, cell type annotation and spatial cell-interaction analysis, yet relatively few openly available datasets contain transcriptomic, proteomic and histological information acquired from the same tissue section. By releasing the underlying Xenium transcripts, cell boundaries, COMET images, protein measurements, and aligned H&E images from multiple human tissues, together with the accompanying visualisation software UnumLocalia, we aim to provide a resource that can support development and benchmarking of future computational approaches. UnumLocalia also enables extraction of integrated single-cell measurements from either the supplied Xenium segmentation or user-defined cell boundaries, allowing identical multimodal datasets to be analysed using alternative segmentation approaches.

Several technical considerations emerged during development of the workflow. Successful completion of sequential Xenium, COMET and H&E staining depended on preservation of tissue morphology throughout repeated cycles of staining, imaging, washing and antigen retrieval. While only minor tissue loss was noted in our samples, tissue robustness is likely to vary considerably between tissue types and fixation conditions, and optimisation may therefore be required for each sample type. We also observed that the rabbit cell segmentation antibodies used during the Xenium workflow appeared to persist despite antigen retrieval, reducing compatibility with some COMET rabbit primary antibodies. Consequently, these antibodies were excluded from the final COMET panel and subsequent analysis. Future workflows may benefit from incorporating additional steps following Xenium processing to remove residual bound antibodies, or excluding rabbit antibodies during the earliest COMET cycles when residual Xenium signal is more pronounced.

Other technical limitations remain common to fluorescence imaging generally. Tissue autofluorescence continues to influence signal interpretation in certain channels, particularly within liver tissue, and careful threshold selection remains important during downstream analysis. We also observed occasional evidence of antibody aggregation, potentially due to storage of antibodies within the COMET fluidic system during long imaging runs. While this had limited impact on the datasets presented here, optimisation of antibody handling may further improve image quality for longer experiments.

An important strength of this workflow is the robustness of multimodal image registration. Because both Xenium and COMET incorporate DAPI nuclear staining, image alignment could be performed accurately using Xenium Explorer, producing highly consistent registration between transcriptomic and proteomic datasets. Alignment of H&E images was more challenging as it relied on nuclear morphology alone, without the assistance of fluorescence intensity, yet the resulting registration remained sufficiently accurate to assess integrated cell and tissue architecture.

This study has several limitations that need to be taken into consideration when interpreting this research. Although only four representative tissue cores are provided, they were intentionally selected to demonstrate the applicability of the workflow across biologically distinct tissue architectures, rather than to create a disease-specific atlas. In addition, the protein panel assessed on these tissues consisted of 15 markers, substantially fewer than the number of detected transcripts. Consequently, the presented analyses are intended to demonstrate methodological capability rather than provide comprehensive biological insight. Similarly, default Xenium cell segmentations were used throughout the current analyses, recognising that cell segmentation remains an active area of research that can substantially influence downstream measurements. By releasing the raw transcript coordinates and imaging data alongside processed outputs, users remain free to apply alternative segmentation approaches and evaluate their effectiveness.

In summary, we demonstrate that Xenium spatial transcriptomics, COMET cyclicIF and H&E staining can be performed sequentially on the same FFPE tissue section while maintaining sufficient image quality for downstream multimodal analysis. Beyond establishing a workflow for sequential multimodal imaging, this study provides an integrated experimental, computational and data resource for the spatial biology community. By combining openly available same-section transcriptomic, proteomic and histological datasets with UnumLocalia for visualisation and single-cell data extraction, we anticipate this resource will facilitate development of computational methods, improve reproducibility across multimodal spatial analyses and accelerate adoption of integrated spatial technologies.

## Figure legends

**Supplementary Figure 1:** Tissues survive multimodal image processing. Images of slide post-Xenium, prior-COMET, and post-H&E (left-right).

**Supplementary Figure 2:** Multimodal integration produces complementary data to identify single cells in tonsil and HCA samples. Single cells were assessed for protein and gene expression in A-C) tonsil and B-F) hepatocellular adenoma (HCA) tissue. Top row shows low-dimensional embeddings computed from (A, D) protein expression, (B, E) gene expression, and (C, F) combined gene and protein features. Middle row shows spatial maps of cluster assignments (from top-row). Bottom row shows marker heatmaps of the protein-only and gene-only clustering results. Bottom row right-inset shows mapping of cluster identities across modalities (Sankey) and quantitative agreement metrics (ARI, NMI). ARI, Adjusted Rand Index; NMI, Normalised Mutual Information.

## Acknowledgements

This work was funded by the Joan Krefft bequest to the A.W. Morrow Gastroenterology and Liver Centre (Royal Prince Alfred Hospital) and Tour de Cure (Application ID RSP-362-2024). The authors gratefully acknowledge Sydney Cytometry and thank the support staff in this core facility for their assistance. They also thank the Charles Perkins Centre Histology Facility for their assistance with this work.

---

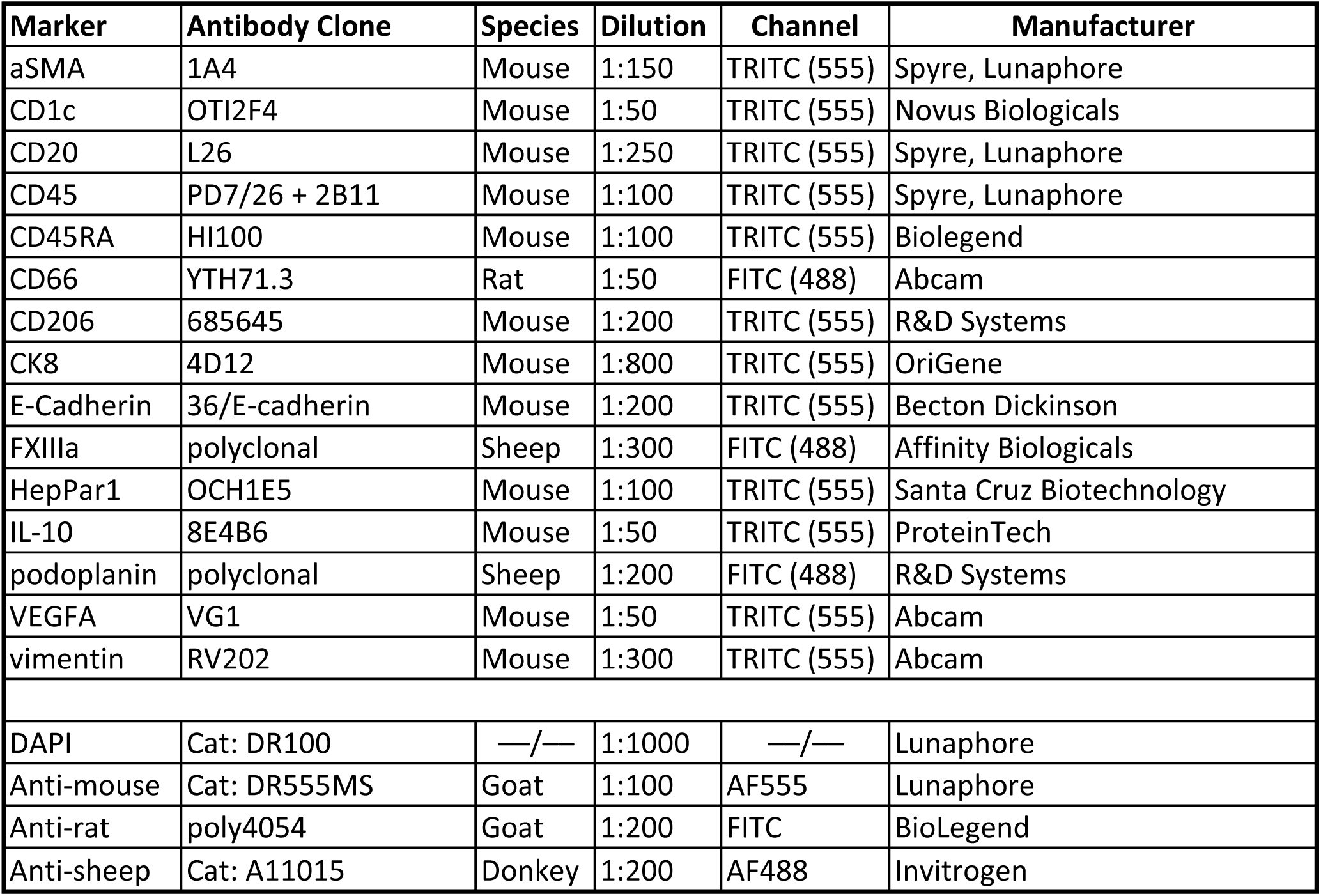

