## Supplementary figures and images for "An open multimodal spatial resource integrating same-tissue transcriptomics, proteomics, and histology"

### SupplementaryFigure1

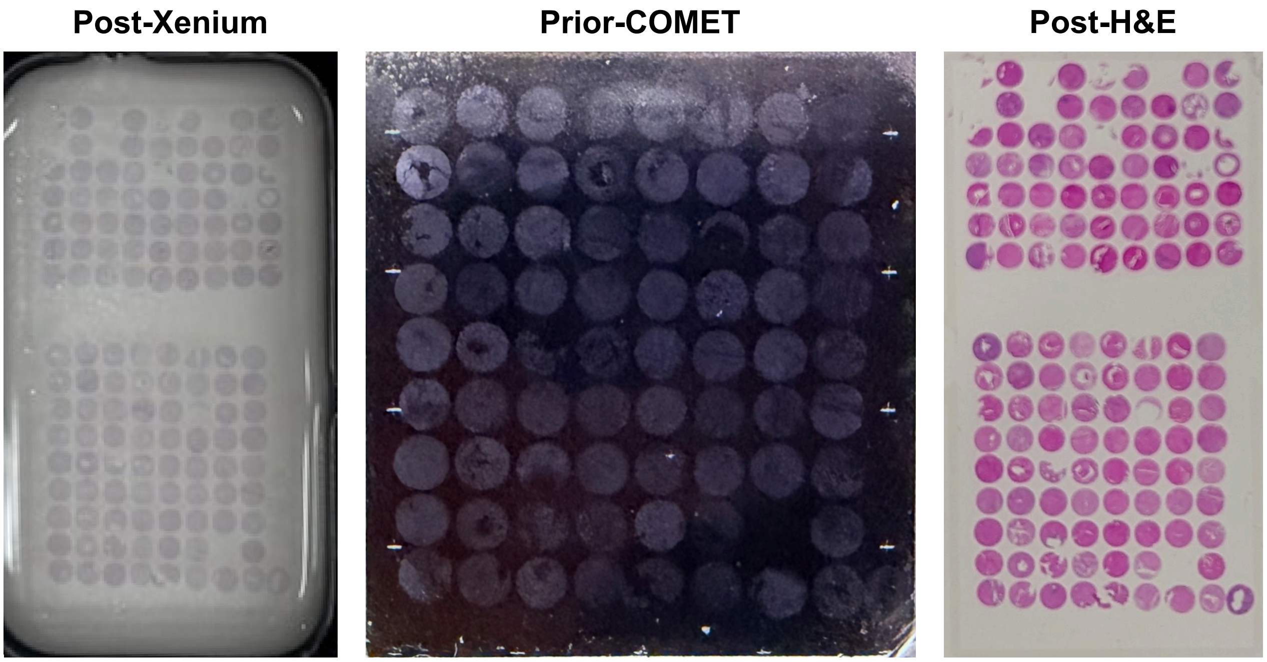
