## SupplementaryFigure2 for "An open multimodal spatial resource integrating same-tissue transcriptomics, proteomics, and histology"

### Tonsil

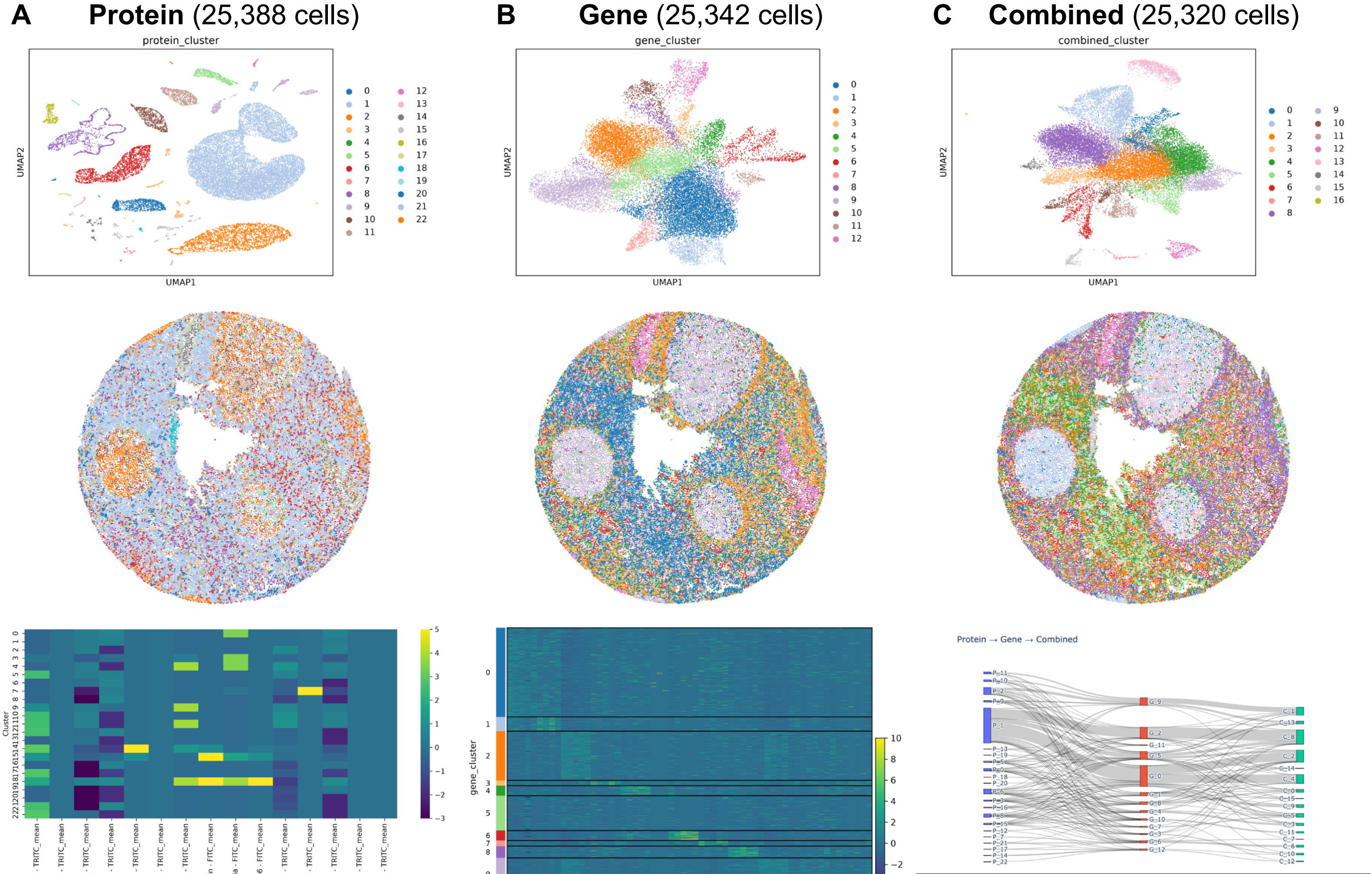


| | Gene vs Protein | Gene vs Combined | Protein vs Combined |
|---|---|---|---|
| ARI | 0.110688844 | 0.446108946 | 0.181697043 |
| NMI | 0.184295962 | 0.603316927 | 0.336822676 |

### HCA

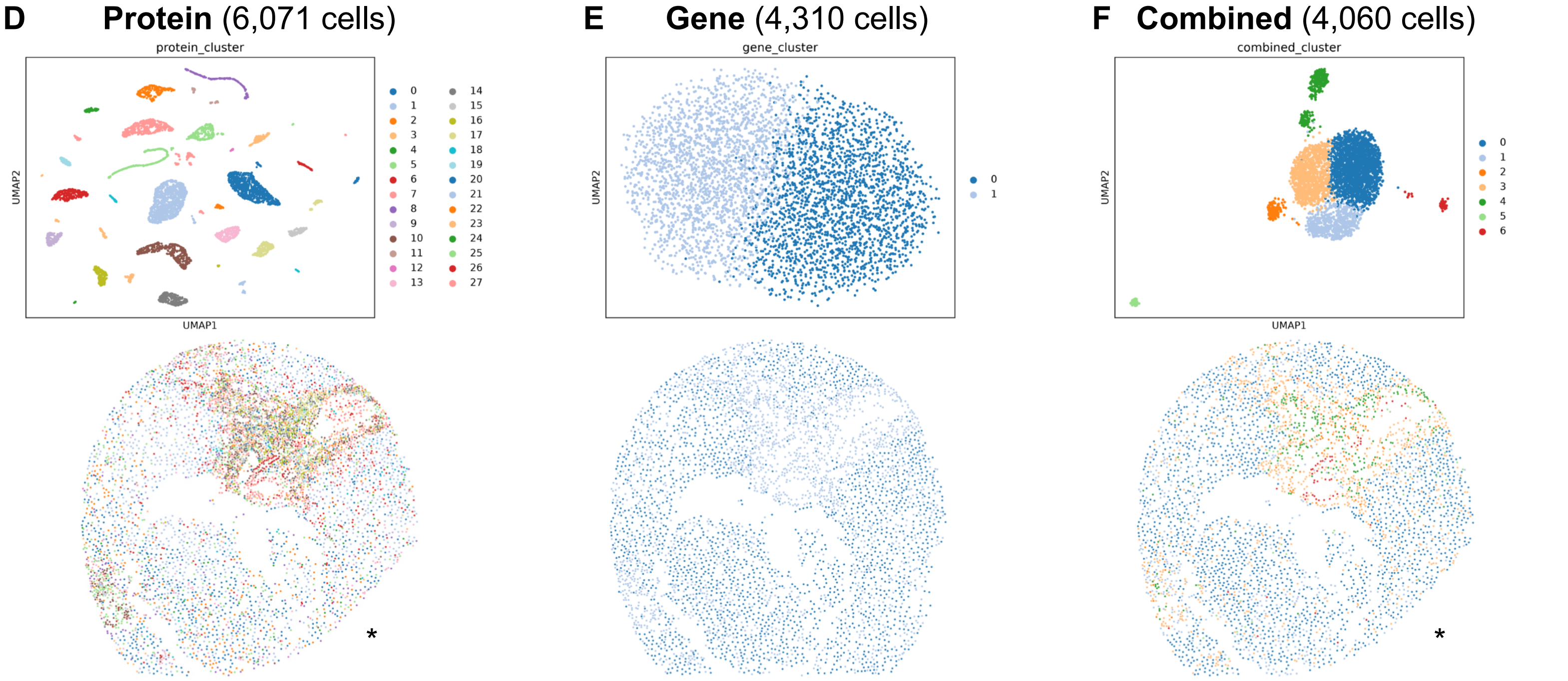


*some cells cut-off during COMET imaging

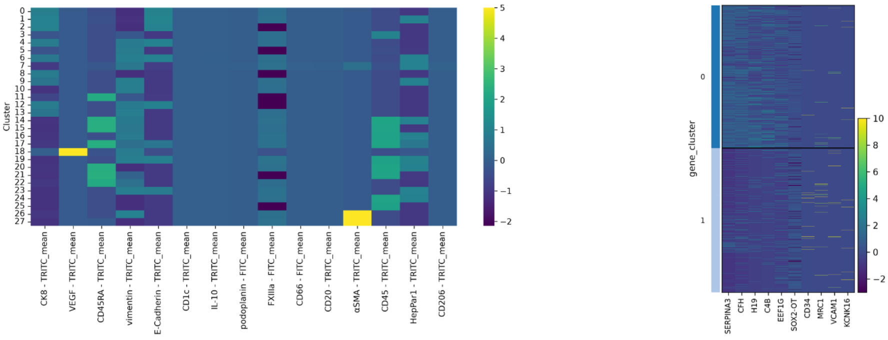


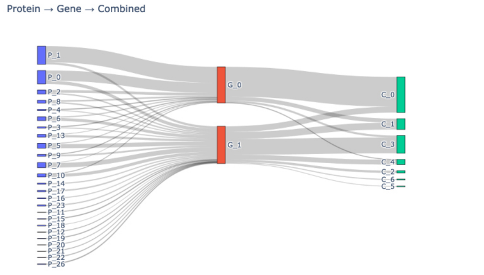


| | Gene vs Protein | Gene vs Combined | Protein vs Combined |
|---|---|---|---|
| ARI | 0.103921433 | 0.291649484 | 0.385720906 |
| NMI | 0.145742222 | 0.253544644 | 0.553559892 |
